# Atlas-CNV: a validated approach to call Single-Exon CNVs in the eMERGESeq gene panel

**DOI:** 10.1101/422337

**Authors:** Theodore Chiang, Xiuping Liu, Tsung-Jung Wu, Jianhong Hu, Fritz J. Sedlazeck, Simon White, Daniel Schaid, Mariza de Andrade, Gail P. Jarvik, David Crosslin, Ian Stanaway, David S. Carrell, John J. Connolly, Hakon Hakonarson, Emily E. Groopman, Ali G. Gharavi, Alexander Fedotov, Weimin Bi, Magalie S. Leduc, David R. Murdock, Yunyun Jiang, Linyan Meng, Christine M. Eng, Shu Wen, Yaping Yang, Donna M. Muzny, Eric Boerwinkle, William Salerno, Eric Venner, Richard A. Gibbs

## Abstract

**Purpose:** To provide a validated method to confidently identify exon-containing copy number variants (CNVs), with a low false discovery rate (FDR), in targeted sequencing data from a clinical laboratory with particular focus on single-exon CNVs.

**Methods:** DNA sequence coverage data are normalized within each sample and subsequently exonic CNVs are identified in a batch of samples (midpool), when the target log_2_ ratio of the sample to the batch median exceeds defined thresholds. The quality of exonic CNV calls is assessed by C-scores (Z-like scores) using thresholds derived from gold standard samples and simulation studies. We integrate an ***ExonQC*** threshold to lower FDR and compare performance with alternate software (VisCap).

**Results:** Thirteen CNVs were used as a truth set to validate Atlas-CNV and compared with VisCap. We demonstrated FDR reduction in validation, simulation and 10,926 eMERGESeq samples without sensitivity loss. Sixty-four multi-exon and 29 single-exon CNVs with high C-scores were assessed by MLPA.

**Conclusions:** Atlas-CNV is validated as a method to identify exonic CNVs in targeted sequencing data generated in the clinical laboratory. The ***ExonQC*** and C-score assignment can reduce FDR (identification of targets with high variance) and improve calling accuracy of single-exon CNVs respectively. We proposed guidelines and criteria to identify high confidence single-exon CNVs.

## Introduction

Copy number variation (CNV) is an important feature of the human genome and can confer disease susceptability^1–4^. The ability to detect CNVs accurately is critical for both genetic diagnostics and to advance understanding their impact on gene function. Next-generation sequencing (NGS) based targeted gene panels are commonly used in clinical genetic testing and various methods^5–13^ have been developed to identify exonic CNVs in gene panel sequence data. Gene panels afford a qualitatively different opportunity to assess small CNVs due to their typically deeper sequence coverage when compared with Whole Genome or Exome Sequencing (WES). Ideally, the methods would detect single-exon CNVs, but this is challenging because a single exon represents one data point, which must exhibit minimal noise and maximal signal compared to multi-exonic CNVs which has corroborating data points. Detecting and reporting CNVs in the clinical context is another challenge as false positives (FPs) must be minimized, while true positives (TPs) are all identified. Achieving the high accuracy required for clinical applications invariably demands validation on alternative platforms, which bears its own set of technical challenges.

Existing CNV tools that have been validated for the clinical setting include VisCap^5^, CoNVaDING^10^, and DeCON^11^, ExomeDepth^12^, and others^6,7,13,14^. CoNVaDING and ExomeDepth are methods which are successful at detecting single-exon CNVs by evaluating individual exon suitability for variant detection and selecting highly correlated samples as reference controls. In these methods, however, each single-exon CNV is treated equally and there is no mechanism to evaluate the confidence of the call. VisCap has relied on human visual scoring to reduce FPs and evaluate small CNVs.

The Electronic Medical Records and Genomics (eMERGE) Network provides an opportunity to develop tools addressing these challenges. Briefly, eMERGE Phase III is the continuation of a NIH program which aims to incorporate genomic information into medical records (25,000 participants) by identifying rare genetic variants using eMERGESeq (targeted gene panel), and their effects in 109 clinically relevant genes, including the ACMG^15^ 56 medically actionable genes. Here, we present Atlas-CNV, a method to identify CNVs even at the single exon level based on the normalized coverage among samples but constrained to the same capture experiment. We incorporate standard deviation (StDev) thresholds to remove low quality exons and samples for controlling FPs. Atlas-CNV produces graphical gene and exon bar plots to allow for visualization by clinicians and diagnosticians. The leveraging of the C-scoring exons to prioritize high quality single-exon CNVs enables significant reduction of FPs and obviates much of the need for costly expert-based validation and reviewing. Atlas-CNV can analyze NGS targeted gene data from a batch of multiplexed samples in a single probe-hybridization capture experiment undergoing identical conditions, which eliminates the need to sequence control samples. This is defined as a midpool experiment which is typically optimized to contain 45-48 samples. Overall, Atlas-CNV is a fast (<2 min per midpool on a single CPU core) and versatile CNV calling method that can be easily integrated into clinical diagnostic pipelines.

We benchmarked and validated Atlas-CNV on known CNVs previously identified by WES and chip array, demonstrated the exon C-scoring feature through simulation on a subset of eMERGESeq samples (sequenced at the Baylor College of Medicine Human Genome Sequencing Center), and assessed its performance by verifying 64 and 29 multi and single-exon CNVs respectively through MLPA. Currently, while eMERGE reports CNVs spanning two or more contiguous exons, our findings support the feasibility of including single-exon CNVs into medical records with the potential for increased diagnostic yield^16^.

## Materials and Methods

Atlas-CNV is available for public download at http://github.com/theodorc/atlas-cnv. with the initial version (0.2) written in Perl (5.12.2) and R (3.1.1). Three inputs are required: (1) GATK DoC interval summary files, (2), a panel design containing target exons, and (3) a sample file with gender and/or midpool groupings.

### Clinical Sequencing

Our clinical pipeline routinely processes about 45 samples in each midpool capture experiment. Briefly, sample DNA is isolated, sheared, ligated to barcode adapters for multiplexing, then incubated with capture probes, and sequenced on Illumina HiSeq 2500 instruments with two midpools loaded on a single flow-cell lane. Paired-end reads are aligned to the hg19 reference using bwa-0.6.2^17^ with GATK-2.5.2^18^ for realignment, recalibration, and depth of coverage calculations (DoC).

### RPKM Normalization and Sample Quality

The read depth (RD) data is normalized at the individual sample level. GATK-DoC output is used to obtain average RD per target, and these values are normalized as a fraction of the sample coverage with RPKM normalization as illustrated in Figure 1A. Essentially, this step converts the average RD per target to the equivalent number of reads (100bp/read) and reports the proportion to the total number of mapped reads in the sample per million. At each exon, the median sample is selected as the reference after removing the 5% outliers (Z-score at 1.96). Then, log_2_ scores of the sample/median ratio are computed accordingly for all exons on all samples as shown in Figure 1B. To assess sample quality, we define a ***SampleQC*** with two components: (1) a 0.2 threshold on the StDev of log_2_ scores for the sample, and (2) a one-way ANOVA F-test at 5% significance applied on the mean RPKM coverage between midpool samples. If either component is not satisfied, the sample is labelled ‘Fail’. The former is determined by computing the StDev at each exon in one theoretical sample containing exactly two allelic copies at every exon in a simple computation where noise was randomly introduced into the sample/median ratio in 5% increments. Assuming a panel of 2000 target exons, we observed that a StDev=0.2 would correspond to approximately 25% variability in the given sample. A midpool with numerous samples failing the ***SampleQC*** would indicate a departure of sample uniformity and possible errors in the workflow (Supplementary Figure S1).

**Figure 1.**
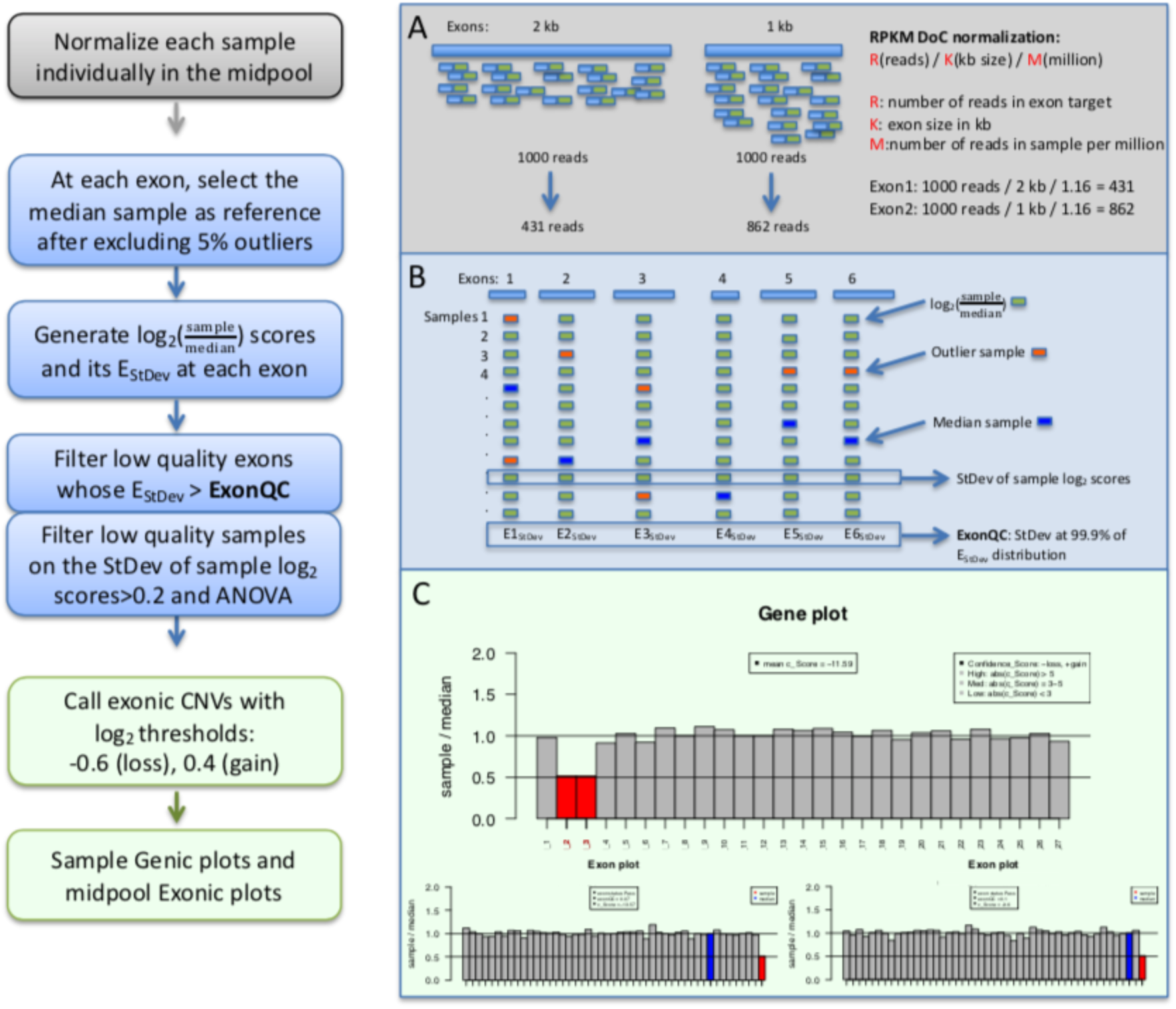
Atlas-CNV method for calling exonic CNVs in a midpool of samples. **(A)** RPKM normalization is first performed on each sample with each exon assigned a single coverage value expressed as a proportion of reads per unit kilobase per sample reads in millions. **(B)** The median sample at each exon is selected as the reference after the 5% outliers are excluded. Log2 scores of sample/median are computed for each sample at that exon and the StDev of these scores is called the **E_StDev_** for that exon. Low quality exons and samples are filtered and flagged. **(C).** CNVs are called with thresholds at the exon level with visual bar plots at the Sample gene level and at the Batch exon level.

### Exon Quality and Calling exonic CNVs

As illustrated in Figure 1B, we also use log_2_ scores to assess exon quality because having even-exon normalized coverage is essential to reduce FPs. Thus, we remove outliers where coverage exceeds a threshold based upon the StDev of log_2_ scores at a given exon (**E_StDev_**) which is calculated on midpool samples. We evaluate the overall distribution of all E_StDevs_ and establish a data derived threshold termed ***ExonQC*** which we define as the E_StDev_ at 99.9% of the E_StDev_ distribution (z=3.921). Exons are labelled ‘Fail’ if its E_StDev_ exceeds ***ExonQC***. To account for E_StDev_ over-inflation due to true exonic CNVs, we first exclude the 5% outliers (approximately 2 samples), to produce a tighter E_StDev_ value which also allows for samples with identical CNVs (kinship). A midpool with many failed exons may indicate aberrant experimental or systematic biases (Supplementary Figure S2). The typical exon fail rate in a given midpool is 0.5%.

To call CNVs, we apply two thresholds on the log_2_ scores (Figure 1C); (1) a user-configurable hard limit of −0.6 for losses and 0.4 for gains, and (2) a soft threshold derived from the data distribution using a Z-score cutoff at 99% (z=2.576), intended as a boundary to threshold calls at the distribution tail. Any log_2_ score exceeding both thresholds are called CNVs. Autosomes and sex chromosomes are analyzed separately using a sample file to define the midpool or gender sub-groupings.

### Visualization and Confidence Score

For visualization, Atlas-CNV produces a sample gene plot with bars representing exons, and also sectioned exon bar plot(s) to display the context of the exonic CNV with all midpool samples. In the latter, the median sample is designated as blue, the test as red, and other samples in grey. An example of a heterozygous CNV loss on 2 contiguous exons in CFTR is shown in Figure 1C.

We define a Confidence Score (C-score) to assign each exonic CNV call (positive and negative scores for duplications and deletions respectively) and suggest three categories: “duplication or deletion”, “*likely* duplication or deletion”, “*uncertain* duplication or deletion” with ranges to denote copy number (Supplementary Material Tables 1-3). C-scores are analogous to Z-scores from the standard normal distribution, where we rescale each exon to unit variance by dividing the individual log_2_ score by the E_StDev_, with an important caveat that assumes the mean log_2_ scores for the given midpool is zero (or diploid). C-scoring standardizes exons on the same comparative scale. For CNVs spanning multiple exons on the same gene, an average of the individual C-scores of the CNV is reported on the gene plot.

### Performance Measures and MLPA

To assess performance, we used samples with CNVs previously identified by WES, and Illumina HumanExome-12v array as our gold standard (GS). We define Sensitivity as the proportion of GS exons (true positives, TP) over the sum of TP and GS exons not called (false negatives, FN); Specificity as the proportion of exons other than GS exons (true negatives, TN) over the sum TN and called exons that are not GS (false positives, FPs); Precision as the proportion of TP over the sum of all positive calls (TP + FP); and False Discovery Rate (FDR) as the proportion of FP over all positive calls. We also define the **‘Reproducibility’** of a tool as the pairwise comparison of two identical runs (A, B) expressed as the proportion of common exons to the union of A and B exons. Finally, an **estimated FDR** (eFDR) is computed per sample using the Robust FDR procedure^19^ which is based on p-values obtained from C-scores.

As our confirmation platform for detected CNVs, we performed MLPA experiments using kits purchased from MRC Holland (www.mlpa.com) for available genes. Samples were processed according to vendor protocols in batches using three controls: NA12878, NA12891, NA12892.

## Results

### Performance Assessment and Comparison

In order to assess Atlas-CNV performance and comparisons to VisCap, we selected 13 clinical samples with a heterozygous gene deletion previously identified by WES and the Illumina HumanExome-12v array as our gold standard (Table 1). Generally, the samples were sequenced in triplicates and divided into technical and biological replicates. The technical replicates use aliquots from the same capture experiment for sequencing while the biological replicates (2 samples: 1-100155, 12-100189) are completely distinct experiments. To compute performance measures, we averaged the replicates of each sample’s comparison to the gold standard and report an overall mean of these 13 samples as bars in Figure 2. We observed high sensitivity and specificity (>99%) across technical and biological replicates, but VisCap has lower precision (80%) and higher FDR (20%) than Atlas-CNV (95%, 4%) which can be attributed to filtering FPs by ***ExonQC***. The legitimacy of filtered FPs was confirmed by their absence in WES data. We also observed that the average E_StDev_ is nearly twice as large in FP calls (E_StDev_ = 0.14 on 45 calls) compared to TP calls (E_StDev_ = 0.079 on 1137 calls).

**Table 1.**
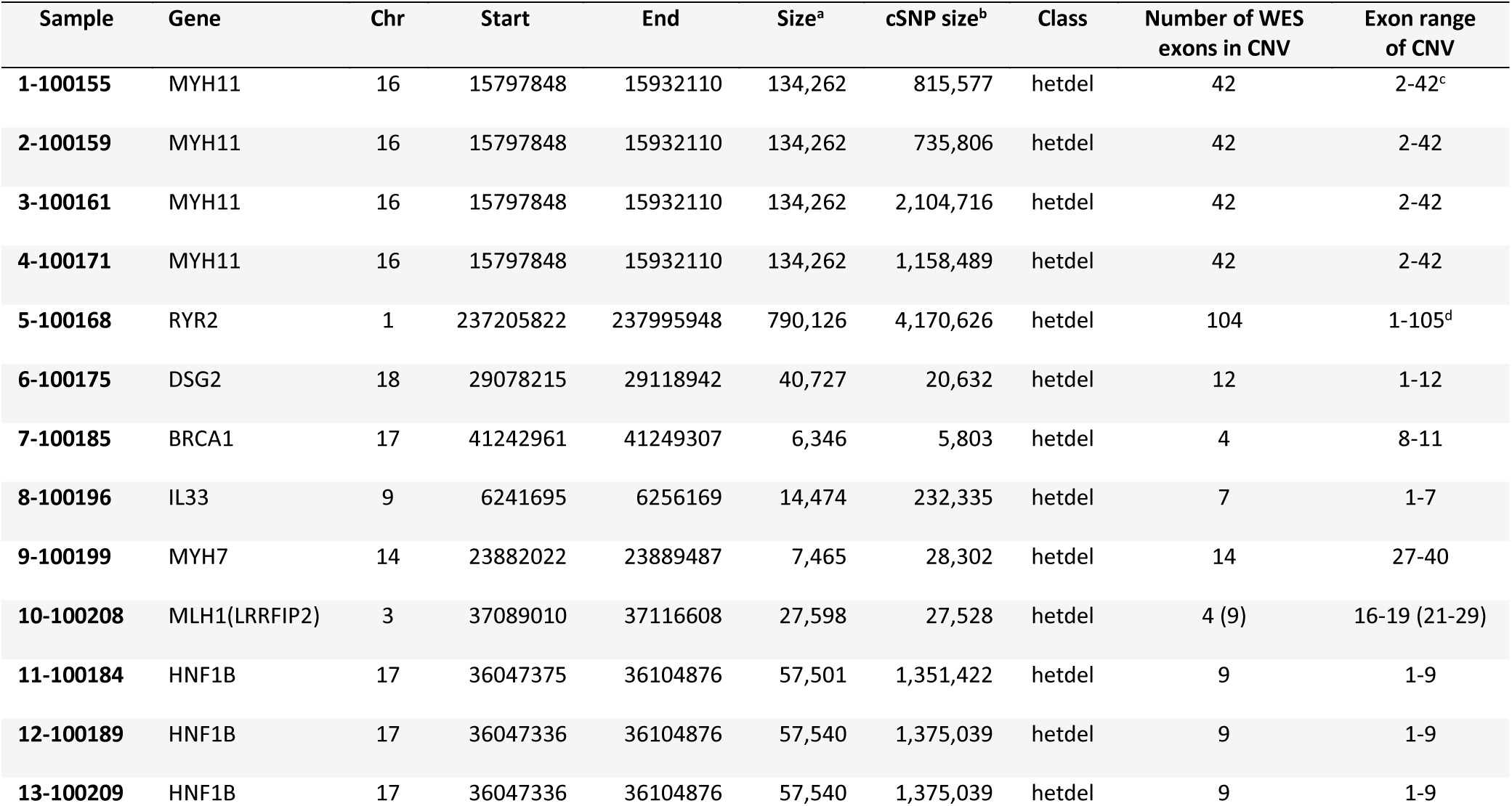
Gold Standard CNVs from 13 clinical samples used to assess Atlas-CNV performance. All CNVs are heterozygous deletions and were previously identified in WES and Illumina HumanExome-12v array with the exception of sample 13 which has no WES. ^a^CNV size from clinical WES. ^b^CNV size from Illumina HumanExome-12v array. ^c^MHY11 exon 42 has 2 targets in WES data. ^d^RYR2 exon 91 is absent in WES.

| Sample | Gene | Chr | Start | End | Size<sup>a</sup> | cSNP size<sup>b</sup> | Class | Number of WES exons in CNV | Exon range of CNV |
| --- | --- | --- | --- | --- | --- | --- | --- | --- | --- |
| **1-100155** | MYH11 | 16 | 15797848 | 15932110 | 134,262 | 815,577 | hetdel | 42 | 2-42<sup>c</sup> |
| **2-100159** | MYH11 | 16 | 15797848 | 15932110 | 134,262 | 735,806 | hetdel | 42 | 2-42 |
| **3-100161** | MYH11 | 16 | 15797848 | 15932110 | 134,262 | 2,104,716 | hetdel | 42 | 2-42 |
| **4-100171** | MYH11 | 16 | 15797848 | 15932110 | 134,262 | 1,158,489 | hetdel | 42 | 2-42 |
| **5-100168** | RYR2 | 1 | 237205822 | 237995948 | 790,126 | 4,170,626 | hetdel | 104 | 1-105<sup>d</sup> |
| **6-100175** | DSG2 | 18 | 29078215 | 29118942 | 40,727 | 20,632 | hetdel | 12 | 1-12 |
| **7-100185** | BRCA1 | 17 | 41242961 | 41249307 | 6,346 | 5,803 | hetdel | 4 | 8-11 |
| **8-100196** | IL33 | 9 | 6241695 | 6256169 | 14,474 | 232,335 | hetdel | 7 | 1-7 |
| **9-100199** | MYH7 | 14 | 23882022 | 23889487 | 7,465 | 28,302 | hetdel | 14 | 27-40 |
| **10-100208** | MLH1(LRRFIP2) | 3 | 37089010 | 37116608 | 27,598 | 27,528 | hetdel | 4 (9) | 16-19 (21-29) |
| **11-100184** | HNF1B | 17 | 36047375 | 36104876 | 57,501 | 1,351,422 | hetdel | 9 | 1-9 |
| **12-100189** | HNF1B | 17 | 36047336 | 36104876 | 57,540 | 1,375,039 | hetdel | 9 | 1-9 |
| **13-100209** | HNF1B | 17 | 36047336 | 36104876 | 57,540 | 1,375,039 | hetdel | 9 | 1-9 |

**Figure 2.**
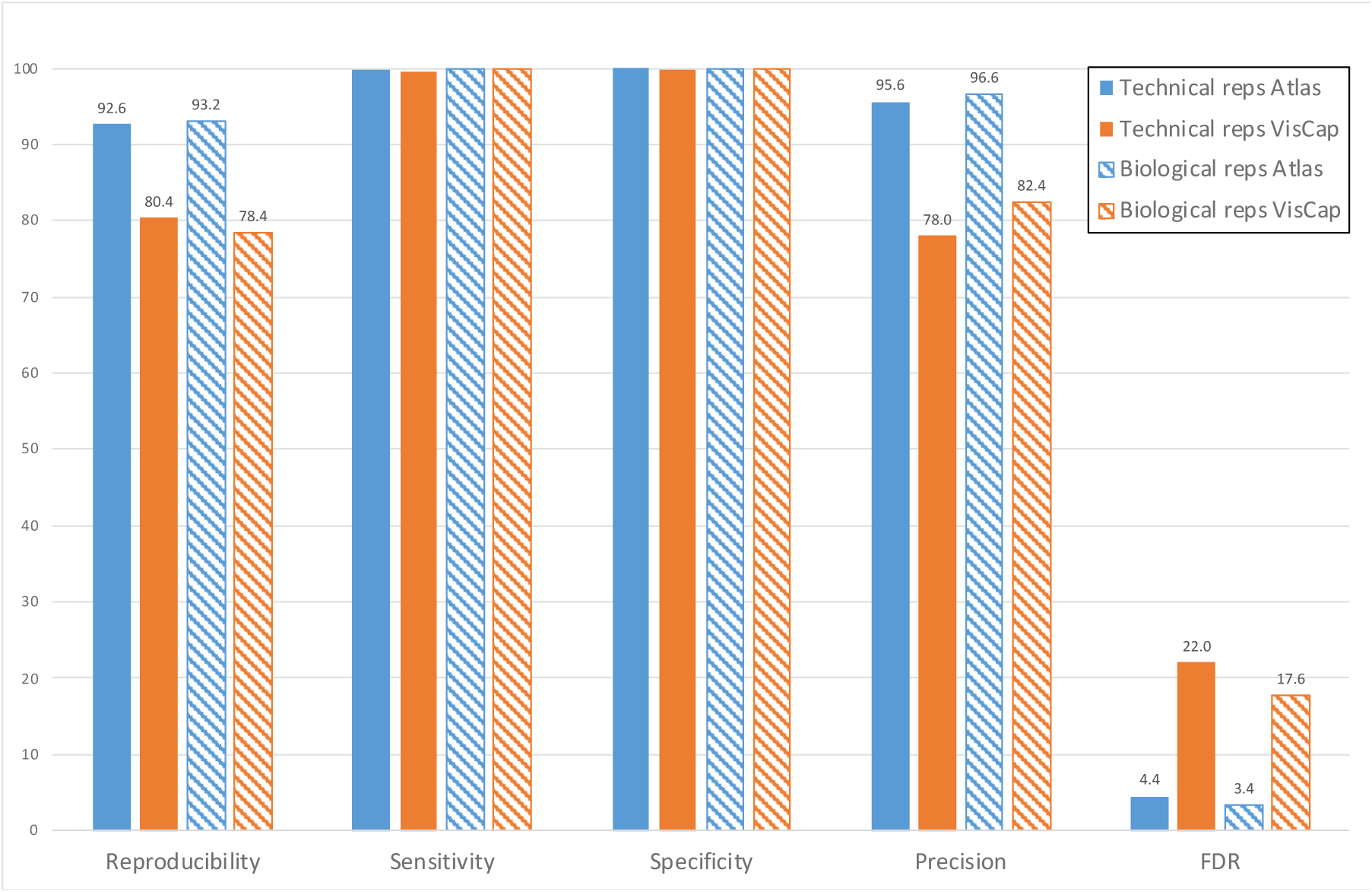
Performance measures comparing Atlas-CNV and VisCap on 13 gold standard samples. Technical replicates were sequenced in triplicates for all samples. Biological replicates were performed on two samples (1 and 12). The performance measure for each sample is first obtained by averaging sample replicates in comparison to the gold standard (GS). Then, an overall mean of the 13 samples are plotted in each bar. Reproducibility is the pairwise comparison of two identical runs (A, B) expressed as the proportion of common exons to the union of exons in A and B. Sensitivity and Specificity are the respective proportions of GS exons (TPs) or exons other than GS (TNs) over the respective sum of these with FNs (failed to call) or FPs (failed to reject). Precision and FDR are the respective proportions of TPs or FPs over all positives.

For reproducibility, pairwise comparisons of replicates were first averaged per sample and then the 13 means were averaged and reported. For the two samples with additional distinct experiments, 3 representative pairwise comparisons were chosen beforehand as the biological replicates (first sample of the technical replicate set), while the remaining 12 comparisons were treated as technical replicates. We report a higher reproducibility in Atlas-CNV (92%) compared to VisCap (80%) indicating ***ExonQC*** may be filtering unrepeated calls in replicate runs.

We estimated an FDR based on p-values from C-scores without prior knowledge of the truth set to confirm the Atlas-CNV FDR of 4%. Using a robust FDR routine under the assumption of a one-sided test^19^, we computed the estimated FDR for the 13 gold standard samples in the range between 0.12 to 14% for technical replicates, and 0 to 15% for biological replicates at p-value cutoffs between 0.009 to 0.01. Although the estimated range is broad, our reported FDR of 4% is within this range of 0-15% and suggests the utility of the procedure on prospective samples without orthogonal confirmation.

### Analysis of eMERGE samples

We analyzed 10,926 eMERGESeq samples from 233 midpools (excluding PMS2^20^ due to highly homologous sequences and 4 midpools with >10% samples failed) with an average of 47 samples per midpool and average coverage of 252x per sample. First, we evaluated the ability of Atlas-CNV and VisCap to call CNVs with at least 2 contiguous exons. Both detected multi-exonic CNVs in 2% of samples with 89% agreement (CNVs identified by both/all CNVs identified) at the sample and gene level (autosomes only). Atlas-CNV and VisCap identified 232 and 184 CNVs respectively with fewer samples failing ***SampleQC*** in Atlas-CNV than VisCap (90 vs 208). Discordant calls (70) were largely made on samples failed by the other tool.

Second, we focused on single-exon detection and initially observed significant discrepancies in the number of these calls. First, Atlas-CNV called nearly 5 times fewer single-exon CNVs than VisCap (2240 vs. 10417; dels=861:5213, dups=1379:5204), and second CoNVaDING, a tool developed for single-exon detection, called even fewer than Atlas-CNV (685; 514 dels, 171 dups). Thus, to reduce the complexities of these comparisons and obtain an estimate of the FDR, we counted the number of single-exon CNVs present in >1% of samples which we assume would likely be artifacts or common CNVs. We report 85% (8818/10417) of VisCap calls, which is observed in only 10 exons, exist in >1% of the samples; 5% (114/2240) for Atlas-CNV (1 exon); and 0% (0/685) for CoNVaDING. This highlights the importance of having a mechanism to automatically filter low quality exons to reduce FPs (present in Atlas-CNV and CoNVaDING but not VisCap). In contrast, if we focus on calls in <1% of the samples (Atlas-CNV:2126, VisCap:1599, CoNVaDING:685) which are more likely to be TPs, our results show 46% (or 741/1599) of VisCap and 42% (or 286/685) of CoNVaDING calls are in common with Atlas-CNV. While the concordance is low, closer examination revealed that missed calls were labelled as multi-exon CNVs in Atlas-CNV or failed to meet the Atlas-CNV passing criteria for either an exon or sample (ExonQC or SampleQC). For example, 58% (or 395/685) of CoNVaDING calls either failed the Atlas-CNV ExonQC (6) or SampleQC (324), were labelled as multi-exon CNVs (47), or flagged as FPs (18), leaving only 4 real missed calls. A similar outcome was observed in the VisCap comparison, but only 15% of calls failed an Atlas-CNV quality control QC (225) or were labelled as multi-exon CNVs (8), leaving 39% of calls as truly missed (or 625/1599). Further examination of the common calls also showed CoNVaDING with a higher mean C-score (7.11 vs 4.9) and lower mean E_StDev_ (0.11 vs 0.14) than VisCap.

Figure 3 summarizes the overall Atlas-CNV analysis. It includes results for PMS2 and X chromosome genes, which the former may require further analyses^20^. 345 samples were identified (172 losses, 173 gains) for multi-exonic CNVs which represents an overall frequency of 3.2% (1.57% losses, 1.58% gains), or 0.03 CNVs/sample. Adding high confidence single-exon CNVs (abs(C-score) >=8) marginally increases the frequency to 4.36% (1.0% losses [109], 0.20% gains [22]). As a relative comparison, the CNV frequency in ExAC^21^ is 1.43%. Interestingly, we detected CNVs in 41 of the 58 ACMG genes (excluding ATP7B) with the highest occurrence in OTC (24) and GLA (24) while CNVs were observed in 38 of 51 non-ACMG eMERGE genes with KCNE1(18) and SLC25A40(11) as top genes.

**Figure 3.**
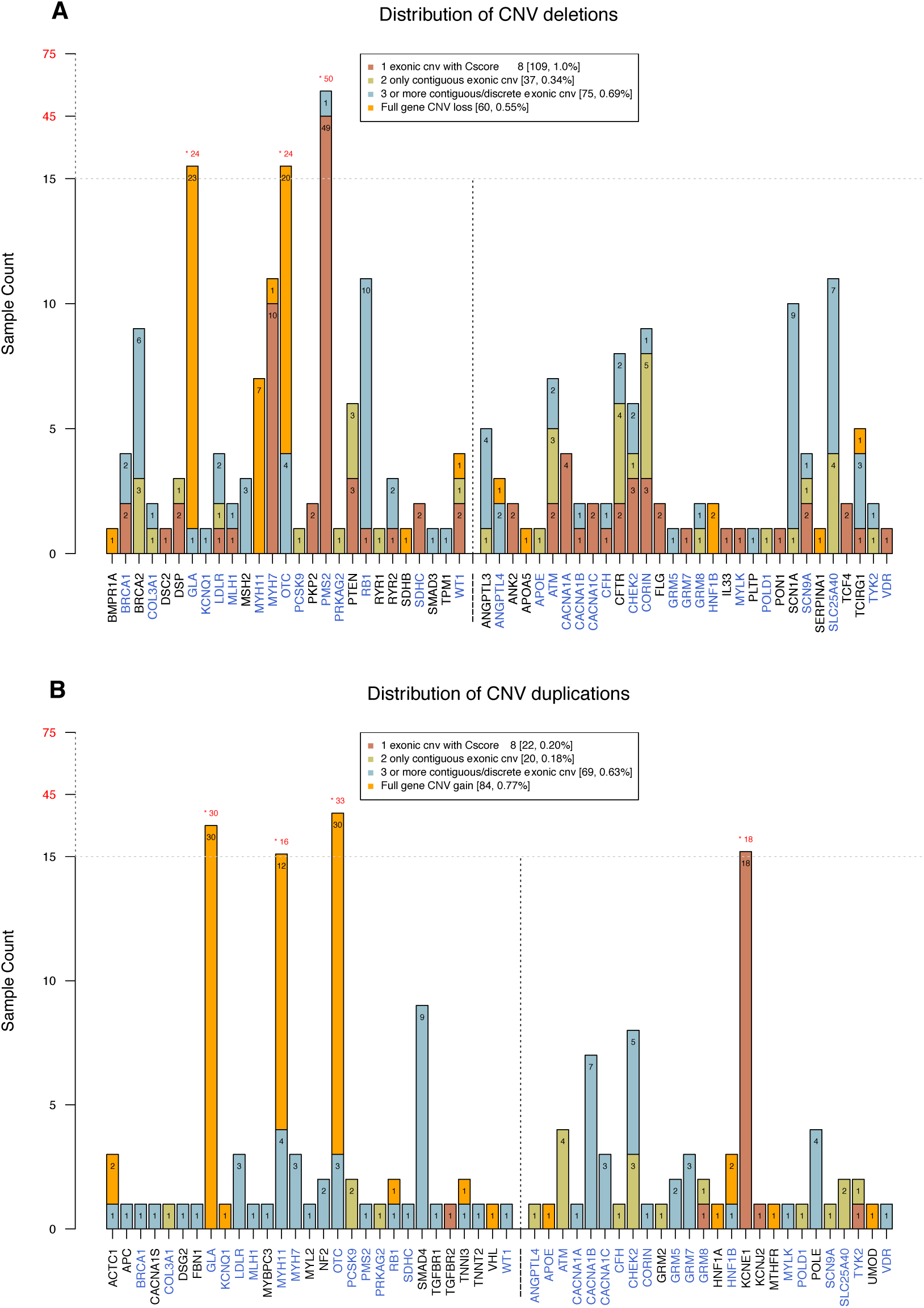
Distribution of CNVs in 10,926 eMERGE samples: deletions (A) and duplications (B). Values in bars represent the sample count broken down by CNV type (single-exon, exactly two exons, three or more, or full gene) with total counts and overall frequency listed in the legend [square brackets]. ACMG genes are grouped alphabetically in the left dotted box and eMERGE specific genes in the right. Gene names labelled in blue were identified in both gains and losses.

### Evaluation of C-score threshold for CNV calling

Across all samples, a total of 2475 exons (on average 11 exons/midpool) were removed by exon filtering with an average E_StDev_=0.36 compared to the 2240 passing exons with a much lower average E_StDev_=0.13. We report an average eFDR of 3.14% across all samples, which coincides with the FDR (4%) reported in the performance assessment study (Figure 2). However, even with low E_StDev_ and FDR, there is still a 10-fold enrichment of single over multi-exon CNVs (2240/232). Therefore, we set out to determine whether C-scores could be used as a secondary assessment of confident of single-exon CNVs primarily on the basis that standalone log_2_ scores do not account for the exon variability whereas C-scores are standardized by this exact variance. First, we computed the expected E_StDev_ in a simulation of log_2_ scores by adding 5% increments of variability into the S/M ratio (Supplementary Figure S3). We determined E_StDev_ at 0.8, 0.13 and 0.16 correspond respectively to 10%, 15% and 20% variability. Therefore, for both eMERGESeq (mean E_StDev_=0.92) and gold standard assessed samples (mean E_StDev_=0.079), the multi-exonic CNVs fall in the 10% noise range. However, eMERGE single-exon CNVs (2240) with nearly twice the mean E_StDev_ (0.14), fall over the >15% noise range (Supplementary Figure S4). Therefore, to control single-exon FPs, both the log_2_ scores and E_StDev_ should be utilized.

We performed a simulation of single-exon deletions to obtain C-score thresholds for optimal sensitivity. Briefly, 100 random samples were chosen from 59% (6,413/10,926) of eMERGESeq samples with no previous single-exon deletion. Each sample was randomly assigned a single-exon deletion by artificially downsizing the read depth by 5% increments from 30% to 50%. The mean C-score and sensitivity were calculated at each coverage increment and plotted on a curve (Supplementary Figure S5). We iterated this analysis using three calling thresholds of −0.4, −0.5, and −0.6 (default), and for all instances in the simulation study, we noted the following: (1) E_StDev_ range 0.08-0.09, (2) eFDR range 3-3.2%, (3) specificity >99%, and most importantly, (4) sensitivity >90% on C-scores >10 where read depth is reduced by >40%. We conclude that these observed ranges for C-score and E_StDev_ are conservative for calling confident single-exon CNVs. When we applied a C-score=>8 and E_StDev_<=0.1 criteria on eMERGE samples, we identified 79 candidate single-exon CNVs (candidates for MLPA validation) which represents a 28-fold decrease from the total of 2240 identified single-exon CNVs. Notably, VisCap and CoNVaDING called 62 and 69 respectively out of these 79.

### MLPA confirmation of CNVs

Sixty-four multi-exon CNVs (34 losses, 30 gains; mean C-score=9.4, E_StDev_=0.087) called by Atlas-CNV were selected for MLPA confirmation (Supplementary Table S4). Although MLPA has its own technical limitations, using it here as the truth set we confirmed 55 CNVs with Atlas-CNV having higher sensitivity (88.8%) and lower FDR (25.0%) compared to VisCap (86.8% and 33.6% respectively). Notably, two confirmed CNVs (Vanderbilt-23 and Columbia-29) were missed by VisCap. The 9 unconfirmed CNVs were compared to the 55 confirmed CNVs and found to have significantly (1) deflated C-scores (5.4, P=8.7e-05), (2) elevated E_StDev_ (0.13, P=2.2e-16), and (3) high CNV genes per sample (>6, P=3.3e-16). A three-prong criteria (C-score >8, E_StDev_ <0.1, CNV genes <3) could easily remove these 9 CNVs and cut the FDR in half to 12.7%. The examination of the actual remaining false positives and negatives indicate missed CNV exons were due to borderline signals on either the gene panel or MLPA platform. Finally, in a further separate analysis, three (28-Northwestern, 30-Columbia, 36-Mayo) samples of the confirmed CNVs (MYH7, LDLR) were also validated using a second gene panel in routine use. Overall, these multi-exon CNV data demonstrate Atlas-CNV performed best with 86% (55/64) confirmed samples and 14% (9/64) failed.

MLPA was also used to evaluate the single-exon CNVs detected by Atlas-CNV (Table 2). Initially, 29 single-exon CNVs were selected for testing from the 79 candidates described above, consisting of 23 high confidence CNVs (22 losses, 1 gain; mean C-score=12.3, E_StDev_=0.081) and 6 borderline confidence CNVs (C-score <8 or E_StDev_ >0.1). However, exact probe reagents for the exon of interest were available for only 14 CNVs (samples 1-14), from which MLPA assays confirmed 10 CNVs (71.4% or 10/14) and was negative for the other 4 (28.6% or 4/14) although 3 of the failed cases had >3 CNV genes per sample. Thus, our overall validation confirmed 90.9% (10/11) of single-exon CNVs (samples 2-11) with C-score >8. One additional single-exon CNV was confirmed from the 15 CNVs (samples 15-29) without exon-specific probes when MLPA assays were carried out using nearby or flanking probes. This results in 14 inconclusive cases for which future investigation is needed using alternate validation methods.

**Table 2.**
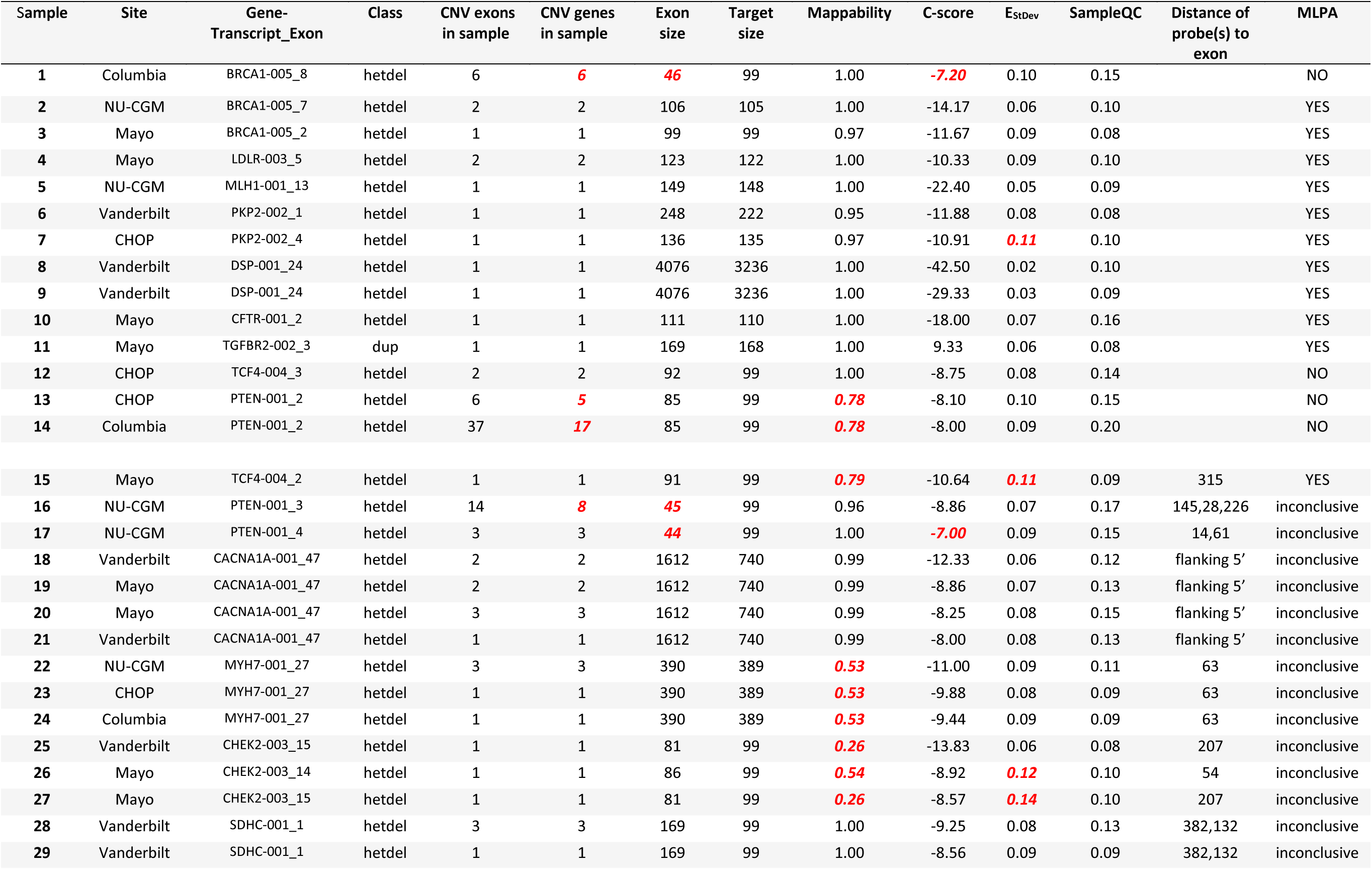
Candidate Single-Exon CNVs from 29 eMERGESeq samples selected for MLPA confirmation. Twenty-three high confidence CNVs (C-score=>8 and E_StDev_<=0.1) and 6 borderline CNVs were tested despite over half (15/29) did not have exact MLPA probes on the exon of interest (see probe distance to exon). High CNV genes per sample, exon size <50bp, or mappability <0.8 may be factors that account for some of the negative and inconclusive cases (see italicized values in bold red). For candidates with C-score >8 and having CNV genes per sample <3, MLPA tests confirmed 90.9% (10/11) of single-exon CNVs.

## Discussion

Prior studies have demonstrated successful CNV detection in DNA sequence data generated from gene panels, but the burden of analyzing single-exon CNVs is a challenge because of high false-positive rates^5,6^. Recently developed tools such as CoNVaDING^10^ and modified versions of ExomeDepth^11,12^, have been designed to identify single-exon CNVs, but lack quality metrics that enable differentiation of different levels of confidence. As a result, clinical laboratories may often ignore single-exon CNVs. Here, we introduced Atlas-CNV, a fast and accurate CNV calling method based on read depth that report confidence scores for each CNV event that are used to reduces FPs. Previously established methods often fail to account for target variability arising from extremes of DNA sequence coverage, while Atlas-CNV implements multiple strategies to cope with these variabilities. Thus, Atlas-CNV’s advantage over other similar methods is the ability to screen or prioritize for high confidence single-exon CNVs. Atlas-CNV also overcomes the limitation of using log_2_ scores as the sole criteria to detect calls. Furthermore, while existing methods^5,10^ have also shown high sensitivity and specificity for detecting single exons in targeted sequence data, our method does not require additional control samples, prior knowledge of model parameters, nor empirical assessments of specific panel designs^11,13^. These necessities render additional calibrations and can significantly increase costs.

Since Atlas-CNV and VisCap are integral components of the eMERGESeq program, we compared the two and our results showed 90% agreement for large or multi-exonic variants. However, given VisCap’s lack of exon filtering, it is difficult to interpret the 39% (or 625/1599) of calls missed by Atlas-CNV for single-exon variants. Furthermore, visual inspection as recommended by VisCap would be a cumbersome task. Therefore, we evaluated the single-exon CNVs with comparison to CoNVaDING calls and conclude Atlas-CNV only missed 4 calls (0.6% or 4/685). We report favorable mean Atlas-CNV C-scores (7.11) and E_StDev_ (0.11) for the 286 common calls even though 57% (164/286) fail the abs(C-score) >8, E_StDev_ <=0.1 criteria. Interestingly, CoNVaDING missed 10 calls that were called by Atlas-CNV (met the abs(C-score) >8, E_StDev_ <=0.1). Thus, this highlights the performance of Atlas-CNV and its ability to prioritize CNV calls to control the FDR and reducing the need for expensive and labor-intensive downstream analysis or assessment.

Our results also show that optimal C-scores>10 can produce >90% sensitivity for single-exon deletions. We therefore propose the following usage guidelines to assist in selecting high confidence CNVs: (1) abs(C-score) >8, (2) E_StDev_ <0.1, (3) exon size >50bp, (4) mappability >0.8, (5) CNV genes <3 per sample. The first two parameters were derived from multi-exonic CNVs; the third is specific to small targets with potential coverage bias, and the last two are considerations to be vetted accordingly since samples with many CNVs could indicate either high FPs or the need for further study. Applying a criterion of only C-score=>8 and E_StDev_ <=0.1, we prioritized 79 significant candidates from the initial 2240 single-exon calls. Out of 11 cases with definitive MLPA assays we confirmed 10 with examples in CFTR, MLH1, and other genes like PKP2 and DSP in which known mutations can increase the risk of arrhythmogenic right ventricular cardiomyopathy, a leading cause of sudden heart failure in young people. Additional frameshift indels were also discovered in the DSP samples, raising the importance of the role single-exon deletions may have for candidate compound heterozygotes in autosomal recessive disorders.

This work advances the confident identification of exonic CNVs, especially in clinical programs deployed at-scale. The application of the Atlas-CNV approach to calling single-exon variants should improve current variant calling standards. We also expect that our knowledge of disease genes will increase as new single-exon CNVs are uncovered, catalogued in public databases and reliably reported. Clinical sites receiving such reports will also benefit patients to obtain better diagnosis and treatment. In conclusion, we have demonstrated Atlas-CNV as a validated approach for clinical laboratories to screen the full spectrum of exonic CNVs in gene panels, with particular focus on single-exon CNVs.

## Acknowledgements

The eMERGE Network phase III work was funded through the following grants: U01HG8657 (Kaiser Permanente Washington, formerly Group Health Cooperative/University of Washington, Seattle); U01HG8685 (Brigham and Women’s Hospital); U01HG8672 (Vanderbilt University Medical Center); U01HG8666 (Cincinnati Children’s Hospital Medical Center); U01HG6379 (Mayo Clinic); U01HG8679 (Geisinger Clinic); U01HG8680 (Columbia University Health Sciences); U01HG8684 (Children’s Hospital of Philadelphia); U01HG8673 (Northwestern University); U01HG8701 (Vanderbilt University Medical Center serving as the Coordinating Center); U01HG8676 (Partners Healthcare/Broad Institute); and U01HG8664 (Baylor College of Medicine).

## Supplementary Material

**Figure S1.**
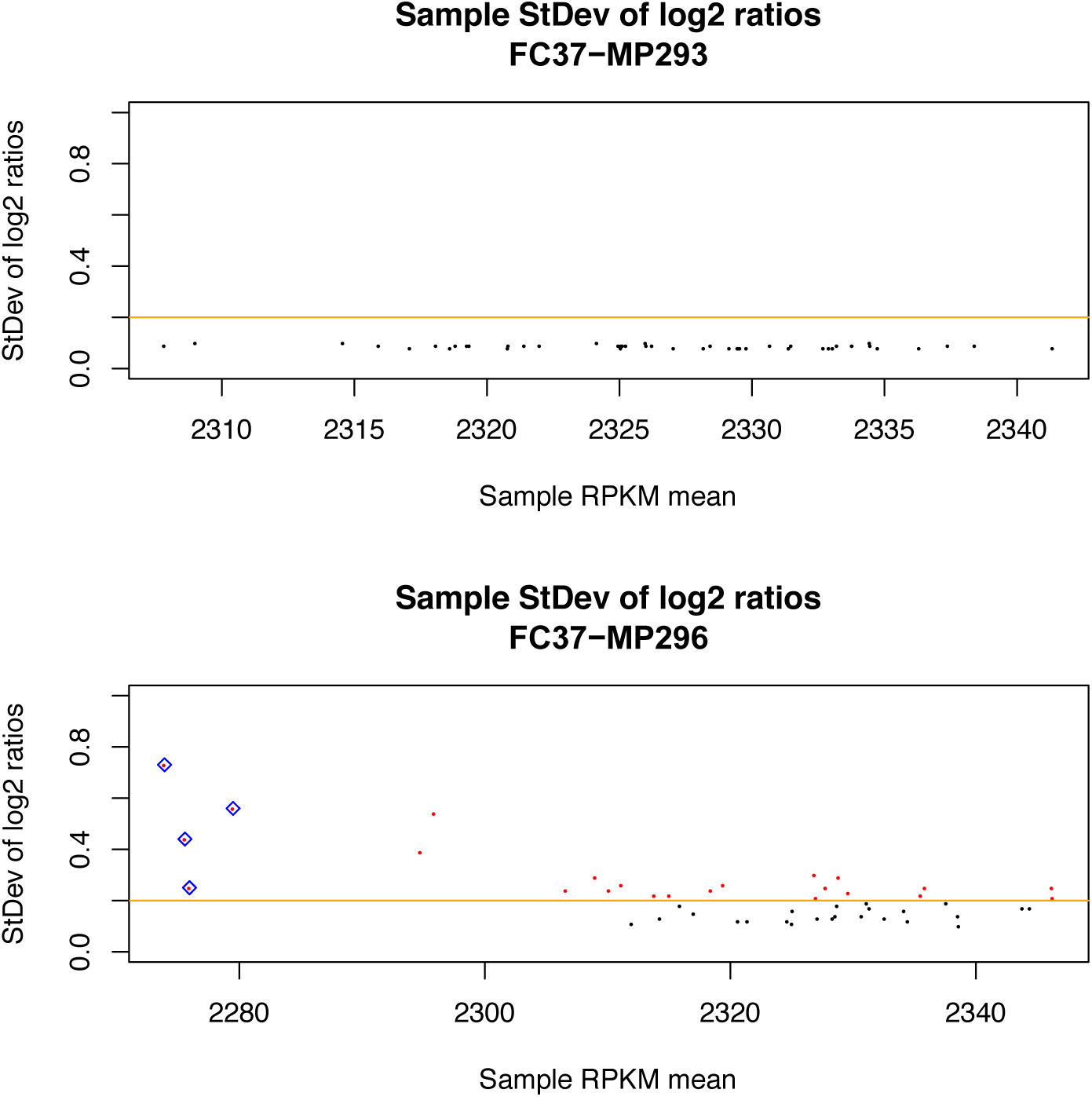
SampleQC. Plots for two midpools (293,296) from a single sequencing flow-cell (37) run containing 47 eMERGESeq samples per midpool. Each data point represents the RPKM mean (x-axis) and the standard deviation of log_2_ ratios (y-axis) of a single sample. The top panel shows midpool 293 where samples have consistent standard deviations below the 0.2 threshold (orange line) whereas the bottom panel show midpool 296 with inconsistencies in sample qualities. Failed samples are labeled in red, and samples with blue diamonds also failed the ANOVA F-test based on their coverage means. The ANOVA test is based solely on the coverage means and will likely be less stringent than the sample standard deviation of log_2_ ratios.

**Figure S2.**
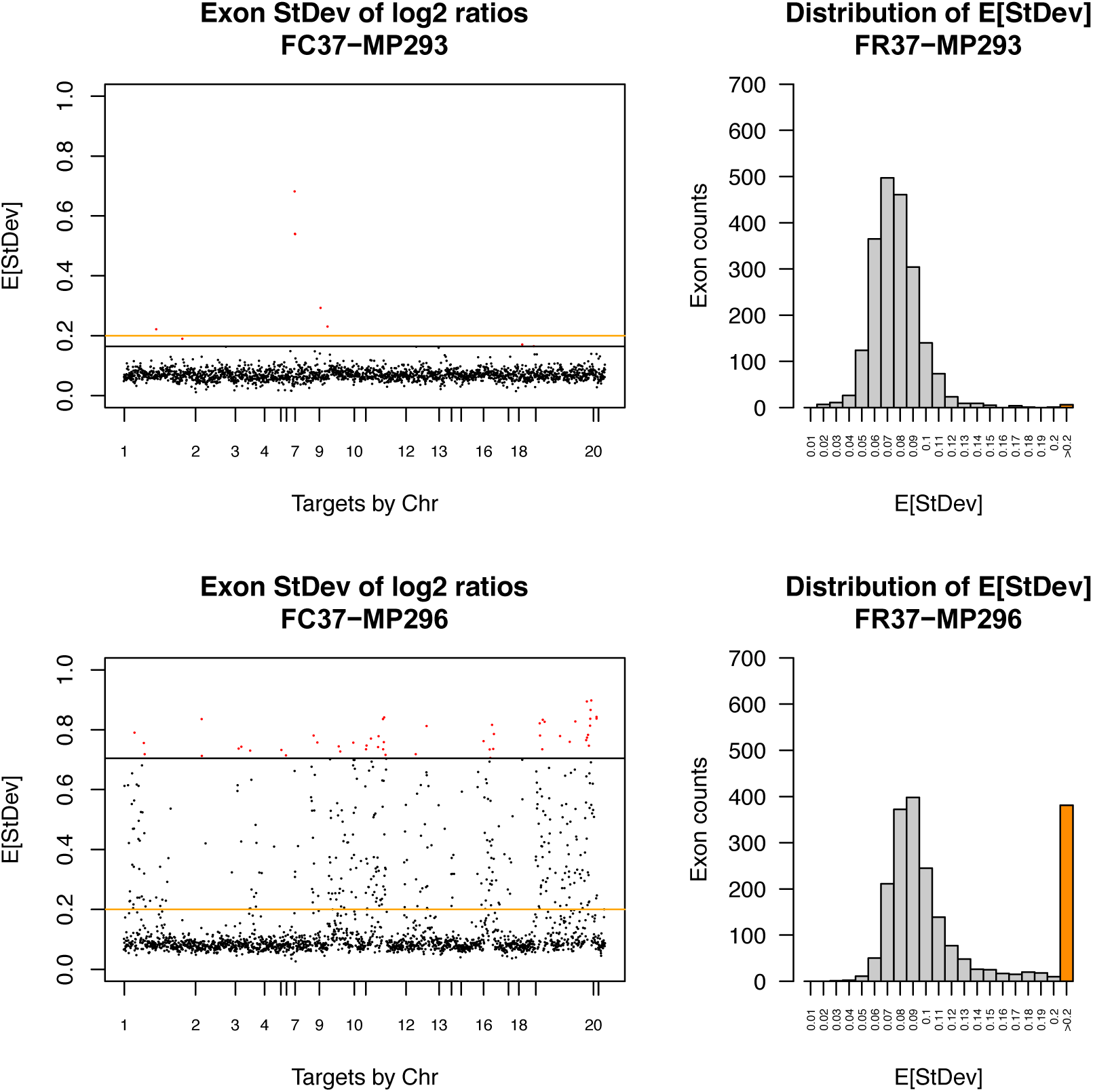
ExonQC. The plots on the left show the Exon standard deviation, E_stDev_ for the same two midpools as Figure 1. E_StDev_ is the standard deviation of log_2_ ratios (sample/median) at each exon on all samples after removing the 5% outliers. The top midpool is typical with only 6 (red) failed exons (6/2066 or 0.3%), whereas the bottom midpool clearly indicate systemic bias (aquarium bubble effect) affecting approximately 18% (381/2066) of exon targets. Orange and black horizontal lines represent a theoretical hard (0.2), or a data derived ExonQC threshold respectively. Histograms on the right are distributions of E_StDev_ which on the bottom midpool illustrates a high count of exons (381) with high variability (E_StDev_>0.2) represented by the orange bar, and a slightly right-shifted distribution. The ExonQC values for these two experiments are 0.16 and 0.70 respectively. Note, the latter value is considerably higher than the theoretical expected value of 0.2 and is a good measure to determine bias in the experiment. The kinds of problematic midpool experiments will undoubtedly affect calling exonic CNVs and inflate the false positives.

## 1 Defining a Confidence Score: *Cscore*

In a Standard Normal distribution, we know the standard *Zscore* are defined as:

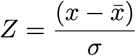

With the same idea, we define a *Cscore* as the following:

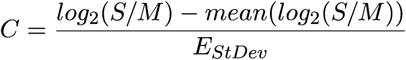

where *S/M* is the exon RPKM ratio of the given sample to the reference median sample, and *E_StDev_* is the standard deviation of *log*_2_(*S/M*) for all the samples within the experiment at a given exon. Note, as most samples are in the diploid state having 2 copies, the mean of *log*_2_(*S/M*) will tend towards 0 essentially because *log*_2_(*S/M*) → 0, and therefore:

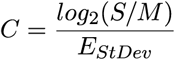

Below, the first two tables show the *Cscores* for Copy Losses and Gains respectively. The values in red are the default thresholds (hard) used in Atlas-pCNV. The third table summarizes the *Cscore* ranges for the types of CNVs, and groups them into ‘High’, ‘Med’, and ‘Low’ categories of confidence.

**Table 1:** Copy Loss Confidence Scores for *E_StDev_*=0.08

| %_Loss | SM_Ratio | log2Ratio | CNV | C_Score |
| --- | --- | --- | --- | --- |
| 0 | 1.00 | 0.00 | diploid | 0.00 |
| 10 | 0.90 | -0.15 | low loss | -1.90 |
| 25 | 0.75 | -0.42 | med loss | -5.19 |
| 34 | 0.66 | -0.60 | threshold loss | -7.49 |
| 40 | 0.60 | -0.74 | 1 copy loss | -9.21 |
| 50 | 0.50 | -1.00 | 1 copy loss | -12.50 |
| 60 | 0.40 | -1.32 | 1 copy loss | -16.52 |
| 90 | 0.10 | -3.32 | 2 copy loss | -41.52 |
| 99 | 0.01 | -6.64 | 2 copy loss | -83.05 |

**Table 2:** Copy Gain Confidence Scores for *E_StDev_*=0.08

| %_Gain | SM_Ratio | log2Ratio | CNV | C_Score |
| --- | --- | --- | --- | --- |
| 0 | 1.00 | 0.00 | diploid | 0.00 |
| 10 | 1.10 | 0.14 | low gain | 1.72 |
| 25 | 1.25 | 0.32 | med gain | 4.02 |
| 32 | 1.32 | 0.40 | threshold gain | 5.01 |
| 40 | 1.40 | 0.49 | 1 copy gain | 6.07 |
| 50 | 1.50 | 0.58 | 1 copy gain | 7.31 |
| 60 | 1.60 | 0.68 | 1 copy gain | 8.48 |
| 90 | 1.90 | 0.93 | 2 copy gain | 11.57 |
| 100 | 2.00 | 1.00 | 2 copy gain | 12.50 |

**Table 3:** Summary of C_Score for CNVs

| Confidence | C_Score | CNV |
| --- | --- | --- |
| High | >5 | duplication |
| Med | 3 to 5 | likely duplication |
| Low | <3 | uncertain duplication (or diploid) |
| High | <-7 | heterozygous deletion |
| High | <-40 | homozygous deletion |
| Med | -3 to -7 | likely heterozygous deletion |
| Low | >-3 | uncertain heterozygous deletion (or diploid) |

**Figure S3.**
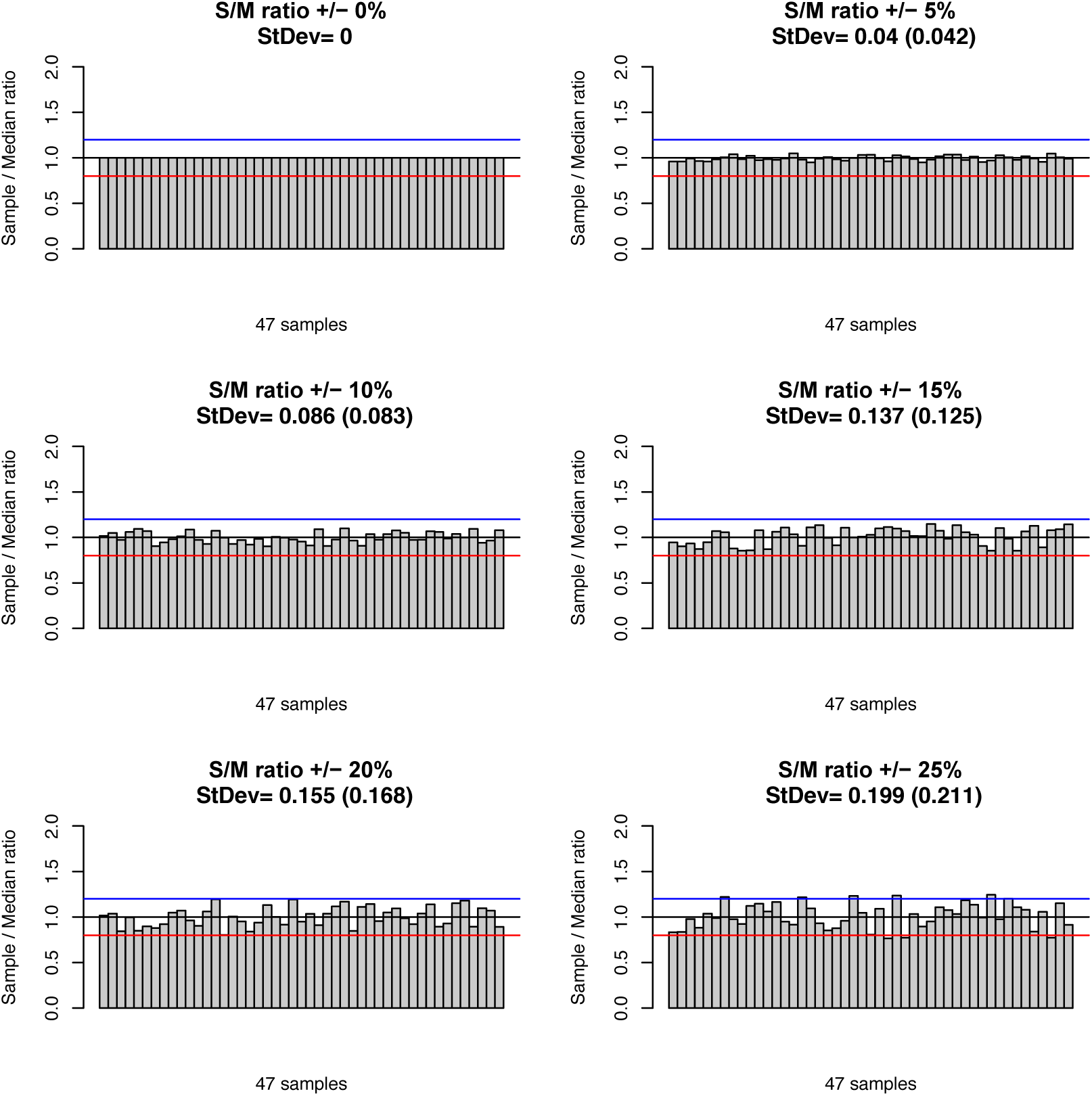
Determining the relationship between E_StDev_ of log_2_ ratios (scores) and % variability in sample to median RPKM ratio (S/M). Six theoretical exon plots on 47 theoretical samples (one midpool) are shown where random noise is added in 5% increments to the **S/M** ratio. The E_StDev_ is reported for the given noise level for one instance and the adjacent value in brackets is the average of 2000 simulated exons instances (akin to eMERGE panel) using the uniform distribution function in R. We conclude the expected upper limit for signal detection should be E_StDev_**=0.2** provided the allowable maximum of 25% noise. Lower values of E_StDev_ will provide higher signal to noise detection abilities. While these values are derived from 47 samples on one exon, analogous strategies can be employed to compute limits for one diploid sample across all ~2000 exons. We employ this strategy for the **SampleQC** (Figure 1) and apply the 0.2 hard threshold to determine whether a sample passes or fails. Blue and red lines denote the 20% boundaries respectively (ie. 1.2 and 0.8). The black line represents the diploid state at 1.0.

**Figure S4A.**
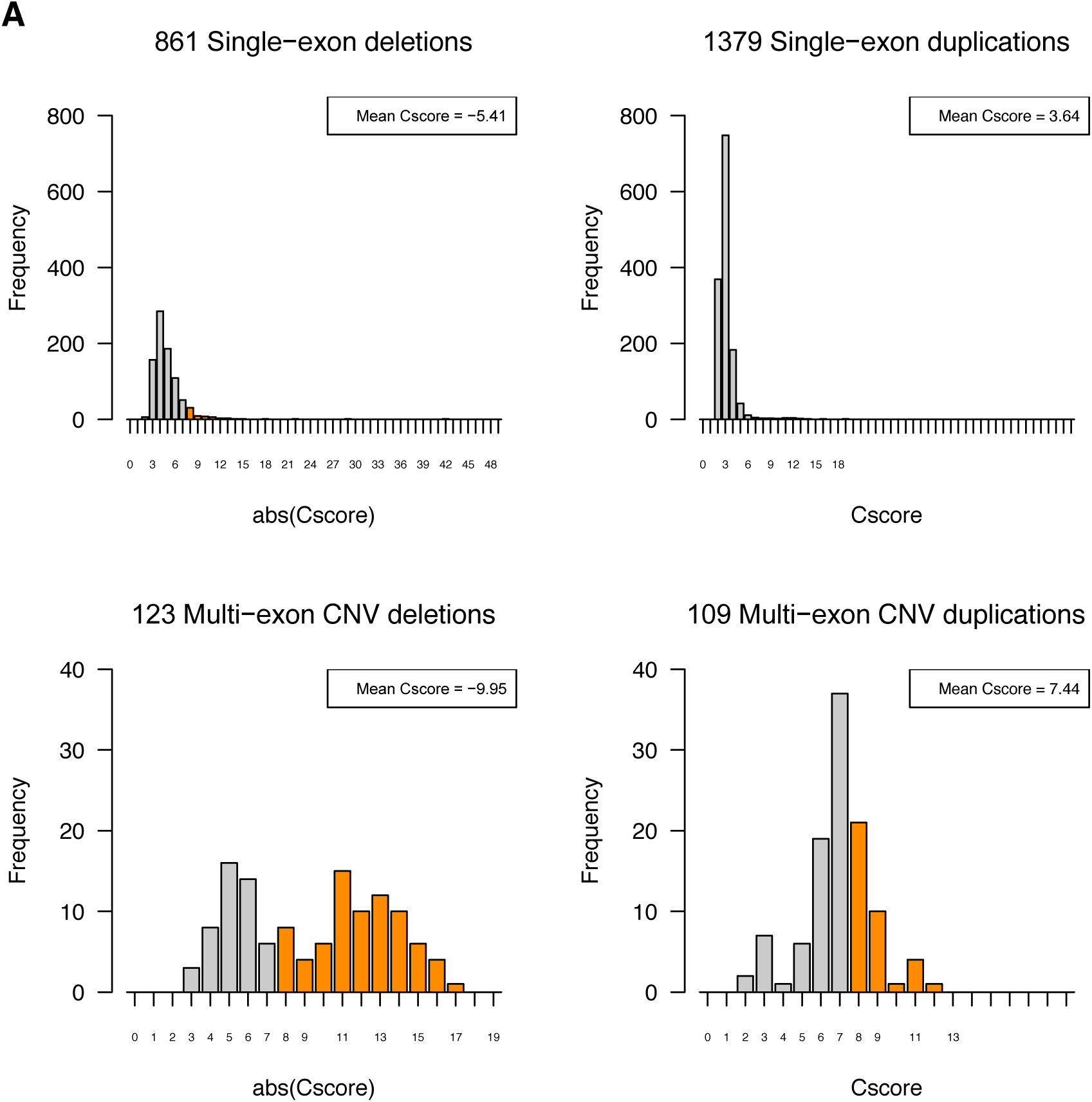
Distribution of C-scores in 10,926 eMERGE samples for single and multi-exonic CNVs excluding PMS2 CNVs. The top panel shows the distribution of C-scores for the 2240 detected single-exon CNVs: 861 deletions (top left) and 1379 duplications (top right), while the bottom panel are for the 232 detected multi-exon CNVs: 123 deletions (bottom left) and 109 duplications (bottom right). The average C-score (magnitude) for all single-exon and multi-exon CNVs are 4.31 and 8.77 respectively. Orange bars represent CNVs that have C-score > 8 in magnitude.

**Figure S4B.**
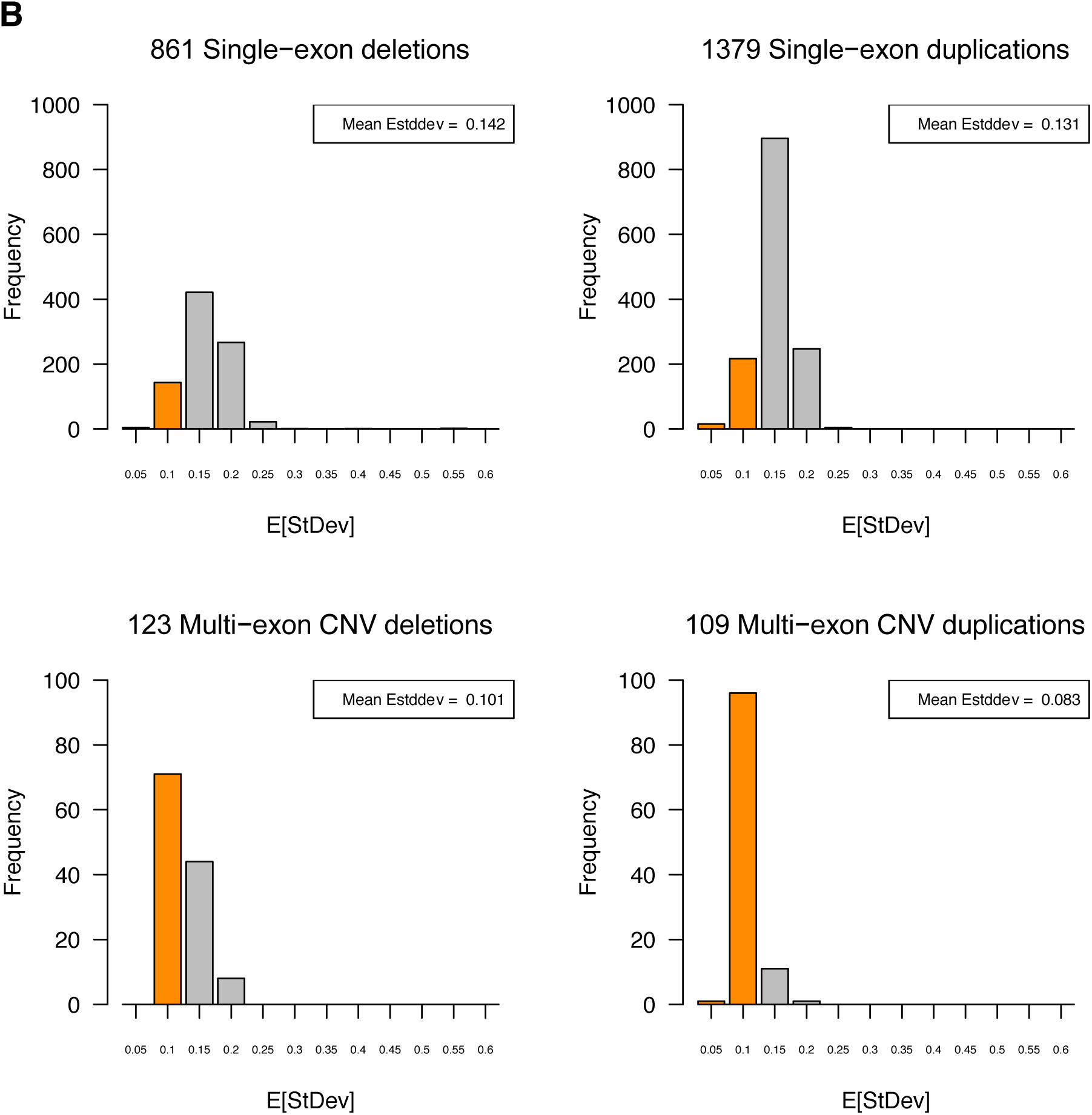
Distribution of E_StDev_ in 10,926 eMERGE samples for single and multi-exonic CNVs excluding PMS2 CNVs. The top panel shows the distribution of E_StDev_ for the 2240 detected single-exon CNVs: 861 deletions (top left) and 1379 duplications (top right), while the bottom panel are for the 232 detected multi-exon CNVs: 123 deletions (bottom left) and 109 duplications (bottom right). The average E_StDev_ for all single-exon and multi-exon CNVs are 0.0923 and 0.135 respectively. Orange bars represent CNVs that have E_StDev_ < 0.1.

**Figure S5.**
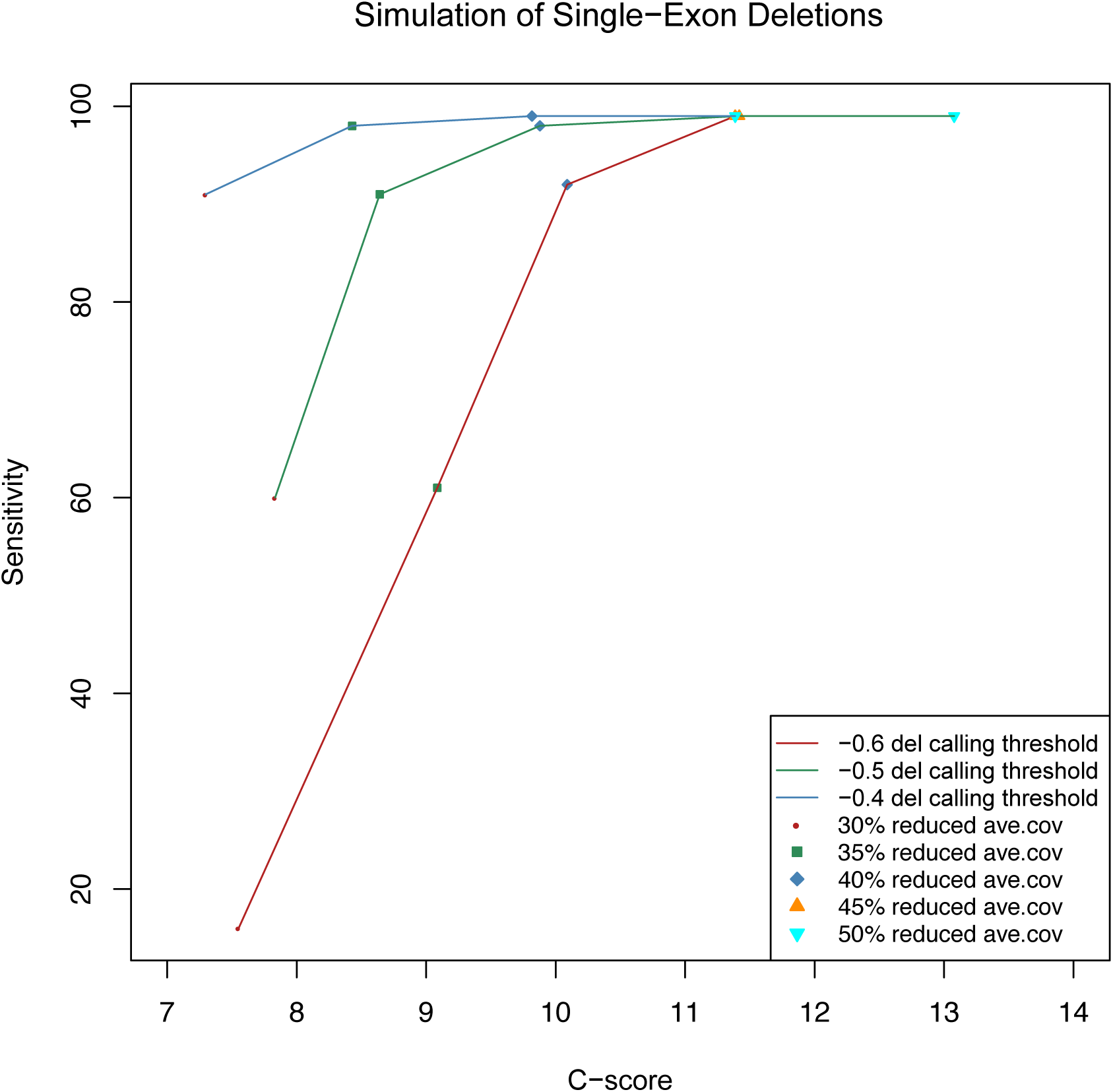
Sensitivity and C-score relationship in a simulation of single-exon deletions in eMERGEsamples. 100 random eMERGE samples with no CNVs were selected and each was assigned one random single-exon deletion. Artificially downsizing the exon coverage was carried out in 5 cycles 30-50% loss and CNVs were called with three log_2_ thresholds (−0.4, −0.5, −0.6). C-scores >10 produced >90% sensitivity.

**Table S4.** *Candidate Multi*-*Exon CNVs from 64 eMERGESeq samples selected for MLPA confirmation*. *The 9 CNVs that failed MLPA confirmation had significantl y* (1) deflated C-scores (5.4, P=8.7e-05), (2) elevated E_StDev_ (0.13, P=2.2e-16), and (3) high counts of CNV genes (>6, P=3.3e-16) when compared to the 55 CNVs confirmed by MLPA.

| Sample | Site | Gene-Transcript_Exons | Class | CNV exons in gene | CNV exons in sample | CNV genes in sample | Mean C-score | Mean $E_{StDev}$ | SampleQC | MLPA |
| --- | --- | --- | --- | --- | --- | --- | --- | --- | --- | --- |
| 1 | NU-CGM | ATM-201_62-63 | dup | 2 | 3 | 2 | 9.87 | 0.07 | 0.09 | yes |
| 2 | Mayo | ATM-201_62-63 | dup | 2 | 2 | 1 | 9.02 | 0.07 | 0.09 | yes |
| 3 | Mayo | ATM-201_62-63 | dup | 2 | 3 | 2 | 9.24 | 0.07 | 0.09 | yes |
| 4 | NU-CGM | ATM-201_2-63 (w/o 28,39,44) | hetdel | 59 | 67 | 3 | -10.14 | 0.07 | 0.15 | yes |
| 5 | Vanderbilt | ATM-201_62-63 | dup | 2 | 2 | 1 | 9.36 | 0.07 | 0.08 | yes |
| 6 | CHOP | BMPR1A-001_3-13 | hetdel | 11 | 11 | 1 | -12.53 | 0.09 | 0.11 | yes |
| 7 | CHOP | CFTR-001_19-21 | hetdel | 3 | 3 | 1 | -14.76 | 0.07 | 0.10 | yes |
| 8 | CHOP | CFTR-001_2-3 | hetdel | 2 | 2 | 1 | -12.73 | 0.08 | 0.09 | yes |
| 9 | CHOP | CHEK2-003_4-16 | dup | 13 | 13 | 1 | 7.12 | 0.09 | 0.09 | yes |
| 10 | CHOP | CHEK2-003_6-16 | dup | 11 | 11 | 1 | 7.25 | 0.08 | 0.10 | yes |
| 11 | Mayo | CHEK2-003_10-11 | hetdel | 2 | 2 | 1 | -16.08 | 0.07 | 0.10 | yes |
| 12 | Vanderbilt | CHEK2-003_9-13 | hetdel | 5 | 5 | 1 | -11.24 | 0.08 | 0.10 | yes |
| 13 | NU-CGM | CHEK2-003_4-14 | dup | 11 | 11 | 1 | 6.50 | 0.10 | 0.10 | yes |
| 14 | NU-CGM | CHEK2-003_4-16 | dup | 11 | 11 | 1 | 6.85 | 0.10 | 0.09 | yes |
| 15 | Mayo | CHEK2-003_4-16 | dup | 13 | 13 | 1 | 5.37 | 0.11 | 0.09 | yes |
| 16 | Mayo | CHEK2-003_2, 4-7, 9-14 | hetdel | 11 | 11 | 1 | -11.78 | 0.10 | 0.12 | yes |
| 17 | Mayo | CHEK2-003_4-5 | dup | 2 | 3 | 2 | 9.52 | 0.06 | 0.08 | yes |
| 18 | Vanderbilt | DSP-001_7-8 | hetdel | 2 | 2 | 1 | -14.13 | 0.08 | 0.18 | yes |
| 19 | CHOP | HNF1B-001_1-9 | hetdel | 9 | 9 | 1 | -14.15 | 0.07 | 0.11 | yes |
| 20 | CHOP | HNF1B-001_1-9 | dup | 9 | 9 | 1 | 8.47 | 0.07 | 0.09 | yes |
| 21 | Vanderbilt | HNF1B-001_1-8 | dup | 8 | 8 | 1 | 8.64 | 0.07 | 0.09 | yes |
| 22 | CHOP | HNF1B-001_1-9 | hetdel | 9 | 9 | 1 | -12.96 | 0.07 | 0.11 | yes |
| 23 | Vanderbilt | HNF1B-001_1-9 | dup | 9 | 9 | 1 | 9.17 | 0.07 | 0.14 | yes |
| 24 | Mayo | KCNQ1-001_3-6 | hetdel | 4 | 5 | 2 | -11.77 | 0.08 | 0.10 | yes |
| 25 | CHOP | KCNQ1-001_1-16, 002_1 | dup | 17 | 17 | 1 | 5.76 | 0.10 | 0.10 | yes |
| 26 | Vanderbilt | LDLR-003_17-18 | hetdel | 2 | 2 | 1 | -9.46 | 0.09 | 0.08 | yes |
| 27 | Vanderbilt | LDLR-003_16-18 | dup | 3 | 3 | 1 | 7.27 | 0.08 | 0.09 | yes |
| 28 | NU-CGM | LDLR-003_13-15 | hetdel | 3 | 3 | 1 | -15.52 | 0.07 | 0.10 | yes |
| 29 | Columbia | LDLR-003_7-8_12_15-16_18 | dup | 6 | 26 | 7 | 4.51 | 0.09 | 0.18 | yes |
| 30 | Columbia | LDLR-003_2-6 | hetdel | 5 | 5 | 1 | -15.10 | 0.07 | 0.09 | yes |
| 31 | CHOP | LDLR-003_13-17 | dup | 5 | 5 | 1 | 7.36 | 0.07 | 0.08 | yes |
| 32 | Vanderbilt | MLH1-001_1ab-2 | hetdel | 2 | 4 | 2 | -13.65 | 0.07 | 0.09 | yes |
| 33 | Vanderbilt | MLH1-001_16-19 | dup | 4 | 5 | 2 | 8.39 | 0.07 | 0.10 | yes |
| 34 | Mayo | MSH2-001_1-3 | hetdel | 3 | 3 | 1 | -14.15 | 0.08 | 0.10 | yes |
| 35 | Mayo | MSH2-001_4-6 | hetdel | 3 | 3 | 1 | -11.78 | 0.08 | 0.09 | yes |
| 36 | Mayo | MYH7-001_28-40 | dup | 13 | 13 | 1 | 8.08 | 0.07 | 0.10 | yes |
| 37 | CHOP | MYH7-001_3-40 | hetdel | 38 | 38 | 1 | -13.84 | 0.08 | 0.17 | yes |
| 38 | Mayo | NF2-001_14-16 | dup | 3 | 4 | 1 | 6.61 | 0.09 | 0.09 | yes |
| 39 | **Broad** | PMS2-001_10-12 | hetdel | 3 | 0 | 0 | -19.88 | 0.07 | 0.12 | yes |
| 40 | CHOP | PMS2-001_1-6, 8-11 | dup | 10 | 11 | 2 | 9.45 | 0.07 | 0.10 | yes |
| 41 | CHOP | SDHB-001_1-8 | hetdel | 8 | 9 | 2 | -10.54 | 0.09 | 0.10 | yes |
| 42 | NU-CGM | SDHC-001_1-3,5-6 | dup | 5 | 5 | 1 | 6.26 | 0.10 | 0.08 | yes |
| 43 | Mayo | SMAD4-201_2-3,11-12 | dup | 4 | 5 | 2 | 8.00 | 0.07 | 0.12 | yes |
| 44 | Vanderbilt | SMAD4-201_2,5,11,12 | dup | 4 | 4 | 1 | 8.73 | 0.06 | 0.09 | yes |
| 45 | CHOP | SMAD4-201_2,3,5,12 | dup | 4 | 4 | 1 | 7.93 | 0.06 | 0.09 | yes |
| 46 | CHOP | SMAD4-201_2,3,5,10,12 | dup | 5 | 5 | 1 | 7.20 | 0.07 | 0.09 | yes |
| 47 | CHOP | SMAD4-201_2,3,5,11,12 | dup | 5 | 6 | 2 | 6.72 | 0.08 | 0.09 | yes |
| 48 | Vanderbilt | TNNT2-008_2-5,003_5,6-11,13-16 | dup | 15 | 15 | 1 | 6.75 | 0.09 | 0.11 | yes |
| 49 | NU-CGM | WT1-001_4-5 | hetdel | 2 | 2 | 1 | -11.06 | 0.09 | 0.09 | yes |
| 50 | Mayo | BRCA1-005_12,14-15 | hetdel | 3 | 3 | 1 | -14.93 | 0.08 | 0.09 | yes |
| 51 | Columbia | BRCA1-005_2-13 | hetdel | 12 | 12 | 1 | -12.63 | 0.09 | 0.11 | yes |
| 52 | Vanderbilt | BRCA1-005_2-7 | dup | 6 | 6 | 1 | 6.79 | 0.10 | 0.10 | yes |
| 53 | CHOP | BRCA2-201_4-7 | hetdel | 4 | 4 | 1 | -10.00 | 0.09 | 0.09 | yes |
| 54 | Mayo | BRCA2-201_6-9, 13-14, 16, 22, 27 | hetdel | 9 | 20 | 2 | -9.27 | 0.08 | 0.14 | yes |
| 55 | CHOP | BRCA2-201_4,5 | hetdel | 2 | 9 | 5 | -4.65 | 0.14 | 0.16 | no |
| 56 | CHOP | BRCA2-201_12,13 | hetdel | 2 | 9 | 7 | -4.11 | 0.16 | 0.15 | no |
| 57 | Columbia | PTEN-001_3-4 | hetdel | 2 | 32 | 10 | -6.52 | 0.13 | 0.19 | no |
| 58 | Columbia | PTEN-001_3-4 | hetdel | 2 | 15 | 4 | -7.21 | 0.11 | 0.14 | no |
| 59 | Columbia | PTEN-001_3-4 | hetdel | 2 | 11 | 4 | -6.62 | 0.11 | 0.13 | no |
| 60 | Vanderbilt | RB1-002_6,9,12,15-16,21,24,26 | hetdel | 8 | 27 | 11 | -5.51 | 0.13 | 0.20 | no |
| 61 | Vanderbilt | RB1-002_5,6,10,15,21 | hetdel | 5 | 15 | 7 | -5.43 | 0.13 | 0.17 | no |
| 62 | CHOP | RB1-002_9,15,16 | hetdel | 3 | 5 | 3 | -4.29 | 0.15 | 0.15 | no |
| 63 | CHOP | RB1_002_5,6,9,10,15 | hetdel | 5 | 20 | 10 | -4.19 | 0.16 | 0.18 | no |
| 64 | NU-CGM | RB1-002_1-27 | dup | 27 | 27 | 1 | 6.83 | 0.09 | 0.10 | yes |

## References

1. Nowakowska B. Clinical interpretation of copy number variants in the human genome. J Appl Genet. 2017;58(4):449-457. doi:10.1007/s13353-017-0407-4

2. Zhang F, Gu W, Hurles ME, Lupski JR. Copy Number Variation in Human Health, Disease, and Evolution. Annu Rev Genomics Hum Genet. 2009;10(1):451-481. doi:10.1146/annurev.genom.9.081307.164217

3. Conrad DF, Pinto D, Redon R, et al. Origins and functional impact of copy number variation in the human genome. Nature. 2010;464(7289):704-712. doi:10.1038/nature08516

4. Zarrei M, MacDonald JR, Merico D, Scherer SW. A copy number variation map of the human genome. Nat Rev Genet. 2015;16(3):172-183. doi:10.1038/nrg3871

5. Pugh TJ, Amr SS, Bowser MJ, et al. VisCap: Inference and visualization of germ-line copy-number variants from targeted clinical sequencing data. Genet Med. 2016;18(7):712-719. doi:10.1038/gim.2015.156

6. Feng Y, Chen D, Wang GL, Zhang VW, Wong LJC. Improved molecular diagnosis by the detection of exonic deletions with target gene capture and deep sequencing. Genet Med. 2015;17(2):99-107. doi:10.1038/gim.2014.80

7. Nord AS, Lee M, King M-C, Walsh T. Accurate and exact CNV identification from targeted high-throughput sequence data. BMC Genomics. 2011;12:184. doi:10.1186/1471-2164-12-184

8. Li J, Lupat R, Amarasinghe KC, et al. CONTRA: Copy number analysis for targeted resequencing. Bioinformatics. 2012;28(10):1307-1313. doi:10.1093/bioinformatics/bts146

9. Bellos E, Kumar V, Lin C, et al. cnvCapSeq: detecting copy number variation in long-range targeted resequencing data. Nucleic Acids Res. 2014;42(20):e158. doi:10.1093/nar/gku849

10. Johansson LF, van Dijk F, de Boer EN, et al. CoNVaDING: Single Exon Variation Detection in Targeted NGS Data. Hum Mutat. 2016;37(5):457-464. doi:10.1002/humu.22969

11. Fowler A, Mahamdallie S, Ruark E, et al. Accurate clinical detection of exon copy number variants in a targeted NGS panel using DECoN. Wellcome open Res. 2016;1:20. doi:10.12688/wellcomeopenres.10069.1

12. Ellingford JM, Campbell C, Barton S, et al. Validation of copy number variation analysis for next-generation sequencing diagnostics. Eur J Hum Genet. 2017;25(6):719-724. doi: 10.1038/ejhg.2017.42

13. Kerkhof J, Schenkel LC, Reilly J, et al. Clinical Validation of Copy Number Variant Detection from Targeted Next-Generation Sequencing Panels. J Mol Diagnostics. 2017;19(6):905-920. doi:10.1016/j.jmoldx.2017.07.004

14. Gambin T, Akdemir ZC, Yuan B, et al. Homozygous and hemizygous CNV detection from exome sequencing data in a Mendelian disease cohort. Nucleic Acids Res. 2016;45(4):1633-1648. doi:10.1093/nar/gkw1237

15. Kalia SS, Adelman K, Bale SJ, et al. Recommendations for reporting of secondary findings in clinical exome and genome sequencing, 2016 update. 2017. doi:10.1038/gim.2016.190

16. Connolly JJ, Glessner JT, Almoguera B, et al. Copy number variation analysis in the context of electronic medical records and large-scale genomics consortium efforts. Front Genet. 2014;5(MAR):1-8. doi:10.3389/fgene.2014.00051

17. Li H, Durbin R. Fast and accurate long-read alignment with Burrows-Wheeler transform. 2010;26(5):589-595. doi:10.1093/bioinformatics/btp698

18. McKenna A, Hanna M, Banks E, et al. The Genome Analysis Toolkit: a MapReduce framework for analyzing next-generation DNA sequencing data. Genome Res. 2010;20(9):1297-1303. doi: 10.1101/gr.l07524.110

19. Pounds S, Cheng C. Robust estimation of the false discovery rate. Bioinformatics. 2006;22(16):1979-1987. doi:10.1093/bioinformatics/btl328

20. Li J, Dai H, Feng Y, et al. A Comprehensive Strategy for Accurate Mutation Detection of the Highly Homologous PMS2. J Mol Diagnostics. 2015;17(5):545-553. doi:10.1016/J.JMOLDX.2015.04.001

21. Ruderfer DM, Hamamsy T, Lek M, et al. Patterns of genic intolerance of rare copy number variation in 59,898 human exomes. Nat Genet. 2016. doi:10.1038/ng.3638

